# Transitional alveolar epithelial cells and microenvironmental stiffness synergistically drive fibroblast activation in three-dimensional hydrogel lung models

**DOI:** 10.1101/2022.05.24.493246

**Authors:** Thomas Caracena, Rachel Blomberg, Rukshika S. Hewawasam, David W.H. Riches, Chelsea M. Magin

**Author notes:** Corresponding Author: Chelsea M. Magin, PhD, 2115 N Scranton Street, Suite 3010, Aurora, CO 80045. These authors contributed equally to this work (Thomas Caracena, Rachel Blomberg).

## Abstract

Idiopathic pulmonary fibrosis (IPF) is a devastating lung disease that progressively and irreversibly alters the lung parenchyma, eventually leading to respiratory failure. The study of this disease has been historically challenging due to the myriad of complex processes that contribute to fibrogenesis and the inherent difficulty in accurately recreating the human pulmonary environment *in vitro*. Here, we describe a poly(ethylene glycol) PEG hydrogel-based three-dimensional model for the co-culture of primary murine pulmonary fibroblasts and alveolar epithelial cells that reproduces the micro-architecture, cell placement, and mechanical properties of healthy and fibrotic lung tissue. Co-cultured cells retained normal levels of viability up to at least three weeks and displayed differentiation patterns observed *in vivo* during IPF progression. Interrogation of protein and gene expression within this model showed that myofibroblast activation required both extracellular mechanical cues and the presence of transitional epithelial cells. Differences in gene expression indicated that cellular co-culture induced TGF-β signaling and proliferative gene expression, while microenvironmental stiffness upregulated the expression of genes related to cell-ECM interactions. This biomaterial-based cell culture system serves as a significant step forward in the accurate recapitulation of human lung tissue *in vitro*, and highlights the need to incorporate multiple factors that work together synergistically *in vivo* into models of lung biology of health and disease.

## Introduction

Idiopathic pulmonary fibrosis (IPF) is a debilitating interstitial lung disease marked by progressive stiffening of the pulmonary extracellular matrix (ECM), which impairs gas exchange, reduces compliance, and leads to respiratory failure within 3 to 5 years^1^. The only FDA-approved drug treatments for IPF slow disease progression but do not reverse altered lung mechanics and function^2-4^. Mechanistic studies of the disease indicate that repeated injury to alveolar epithelial type II (ATII) cells induces differentiation into alveolar epithelial type I (ATI) cells as part of normal re-epithelialization. In fibrosis, a population of KRT8+ transitional cells persists^5^. These KRT8+ transitional cells do not fully differentiate into ATI cells and can display a profibrotic phenotype that initiates an aberrant wound-healing response in surrounding fibroblasts. This response is mediated by secreted factors capable of inducing fibroblast migration, proliferation, and activation, such as transforming growth factor-β (TGF-β), platelet-derived growth factor (PDGF), and connective tissue growth factor (CTGF)^6, 7^. Over time these interactions result in pulmonary fibrosis, characterized by fibroblast differentiation into myofibroblasts, excess ECM deposition^8^, and local tissue stiffening. In this way, IPF progresses via a positive feedback loop where an uncontrolled healing process results in widespread ECM remodeling and the remodeled ECM activates nearby fibroblasts^9, 10^. While these results are consistent with previous studies reporting on the influence of ECM stiffness on fibroblasts^10-14^, few studies have investigated the influence of local stiffening on alveolar epithelial cells^12, 15^ and how these responses may also reinforce the progression of fibrosis. Therefore, we engineered a biomaterial-based lung tissue model to enable study of both the cellular crosstalk dynamics within the distal lung and changes in microenvironmental mechanical properties during the initiation of fibrosis.

There is a critical need for improved models of fibrotic disease that incorporate both ATII cells and fibroblasts in a microenvironment that closely replicates lung architecture and mechanics to facilitate the study of fibrosis and the development of new anti-fibrotic treatments. Until recently, much of our understanding of the mechanisms underlying IPF has been gained from studying cells on supra-physiologically stiff tissue culture plastic in two-dimensional (2D) monoculture or in animal models that fail to recapitulate the progressive nature of IPF in humans and make it difficult to isolate specific cell-matrix interactions, a key driver of the disease^7, 16-19^. Advanced three-dimensional (3D) cell culture models that incorporate multiple cell types^20, 21^, physiological substrate mechanical properties^22, 23^, and more relevant geometries^24-26^ are emerging to overcome these limitations. Organoids are one example of an effective tool for the study of cell-cell interactions in multicellular co-cultures as an additional factor in directing cell behavior and creating a better approximation of the native physiological environment^21, 27^. For instance, Tan *et al*. fabricated nascent organoids by embedding primary human bronchial epithelial cells, microvascular lung endothelial cells, and lung fibroblasts within Matrigel matrices^28^ and demonstrated that epithelial injury was sufficient to induce fibroblast activation^21^. This study, while a promising step forward in the recapitulation of the native cellular environment *in vitro*, did not incorporate any control over the extracellular environment and was subject to the extracellular mechanical and chemical influences of the Matrigel cell culture substrate. In a study demonstrating the feasibility of controlling extracellular mechanical cues using decellularized ECM (dECM), Nizamoglu *et al*. recently used a ruthenium-based crosslinker to increase the stiffness of porcine lung ECM-derived hydrogels, showing that this dECM hydrogel exhibited controllable mechanical properties and that stiffening this hydrogel increased lung fibroblast activation^23^. In contrast to these naturally-derived culture platforms, synthetic biomaterials such as poly(ethylene glycol) (PEG) offer enhanced customizability in exchange for a lack of innate bioactivity, providing an inert hydrogel backbone that can be extensively modified to recapitulate specific chemical and mechanical traits. Controlling polymerization parameters allows for geometric control of the resulting hydrogels, as demonstrated by Lewis *et al*. through the use of a photodegradable PEG hydrogel to create a spherical alveolar scaffold for the culture of murine ATII cells^24^. Extending experimental models to encompass control over both the cellular landscape and the extracellular environment could refine fibrosis models by accounting for cell-cell and cell-matrix interactions that contribute to the progression of fibrosis^9, 20, 29^.

Here, we present a strategy for engineering 3D lung models using poly(ethylene glycol)- norbornene (PEG-NB) hydrogels that present precisely defined 3D geometries and mechanical properties that recapitulate healthy or fibrotic lung tissue to allow assessment of cross-talk between fibroblasts and ATII cells. Hydrogel microspheres (median diameter = 171 ± 42 µm) that accurately replicate the shape and size of human alveoli^24, 30^ were produced by emulsion polymerization. Magnetically aggregating these microspheres, along with primary murine ATII cells, formed a larger acinar structure, which was then embedded along with primary murine fibroblasts within a hydrogel that recapitulated the elastic modulus (E) of healthy (E = 2-5 kPa) or fibrotic (E = 10-30 kPa) lung tissue ^22^. These 3D lung models maintained cellular viability throughout the constructs over at least three weeks post embedding. Over that time, ATII cells seeded into these structures began to differentiate into ATI cells, with a significant proportion (20-25%) displaying a transitional cell phenotype marked by expression of KRT8. The proportion of activated fibroblasts was highest in the ATII-fibroblast co-cultures embedded in stiff hydrogels (44.0% ± 7.1) compared to the same co-culture in a soft hydrogel (16.4% ± 2.1) or fibroblast monoculture in the same stiff hydrogel (21.0% ± 1.0). Gene expression analysis suggests that co-culture promotes cellular proliferation pathways, whereas microenvironmental stiffening induces an ECM-remodeling response. These results suggest that the combination of transitional ATII cells and exposure to a stiff microenvironment synergistically contribute to fibroblast activation. These findings highlight the roles played by the lung epithelium and the mechanical microenvironment in the progression of pulmonary fibrosis, demonstrating the value of this novel *in vitro* model that reconstructs cell-cell and cell-matrix interactions observed *in vivo*.

## Experimental

### Fabrication of 3D Lung Models

#### Synthesis of PEG-NB

The terminal residues of eight-arm 10 and 40 kg/mol PEG-hydroxyl macromers (JenKem Technology) were conjugated with norbornene functional groups through a radical-mediated step-growth reaction adapted from an established procedure^31^. Briefly, poly(ethylene glycol)-hydroxyl (PEG-OH; eight-arm; 10 kg/mol or 40 kg/mol; JenKem Technology) was lyophilized. Next, 5 g PEG-OH was dissolved in anhydrous dichloromethane (DCM; Sigma-Aldrich) in a flame-dried Schlenk flask under moisture-free conditions. 4-Dimethylaminopyridine (DMAP, Sigma-Aldrich; 0.24 g, 0.02 mol) was added to the flask. Pyridine (Fisher; 1.61 ml, 0.02 mol) was injected dropwise to the reaction mixture. In a separate flask, N,N’-Dicyclohexylcarbodiimide (DCC, Fisher; 4.127 g, 0.02 mol) was dissolved in anhydrous DCM. Norbornene-2-carboxylic acid (Sigma-Aldrich; 4.9 ml,0.04 mol) was added dropwise to the flask. The mixture was stirred at room temperature for 30 minutes and then filtered through Celite 545 (EMD Millipore). The filtrate was added to the first flask and allowed to react for 48 h in a light-protected environment. Byproducts were removed by mixing with an aqueous solution of sodium bicarbonate (Fisher). The organic phase was concentrated with rotary evaporation and the product was precipitated with diethyl ether (Sigma-Aldrich) overnight at 4°C. The resulting polymer was dialyzed (1 kg/mol MWCO, Repligen) against 3.5 L of deionized water at room temperature. The water was changed four times over 72 h, after which the polymer was lyophilized to obtain a solid white product. End-group functionalization of the resulting PEG-NB and the purity of the product were confirmed by nuclear magnetic resonance (NMR) spectroscopy. The ^1^H NMR spectrum was recorded on a Bruker DPX-400 FT NMR spectrometer (300 MHz). Chemical shifts for protons (^1^H) were recorded as parts per million (ppm) relative to a residual solvent. Only synthesis products with greater than 90% functionalization were used (Figure S1, S2).

#### Emulsion Polymerization of Hydrogel Microspheres

PEG-NB (eight-arm 40 kg/mol) was reacted with an MMP3-degradable dithiol crosslinker (Ac-GCRDGAPFALRLVDRCG-NH2; GenScript)^32^ via emulsion polymerization^33^ by a photoinitiated thiol-ene reaction to create microsphere templates that enable cellular remodeling of the microenvironment^34, 35^. Parallel plate rheology identified a microsphere hydrogel formulation with an elastic modulus in the range of healthy lung tissue (2-5 kPa)^22^.

The PEG-NB (6 wt%) was reacted with the MMP3-degradable crosslinker at a ratio of 0.7 thiol reactive groups to NB reactive groups along with pendant peptide mimics of fibronectin (CGRGDS) and laminin (CGYIGSR; GL Biochem) (1 mM final concentrations). First, all components were dissolved individually in neutral (4-(2-hydroxyethyl)-1-piperazineethanesulfonic acid (HEPES; ThermoFisher) buffer and sonicated at 40°C for 10 minutes to create stock solutions. The PEG-NB and peptide mimics were then combined in a microcentrifuge tube (Tube 1) with additional HEPES to bring the total volume to 1.5 mL. The crosslinker was combined with 15 mM tris(2-carboxyethyl)phosphine (TCEP; Sigma-Aldrich) and thiolated magnetic nanoparticles (MNPs; 0.3 mg/mL final volume), synthesized as previously described^36^ in another microcentrifuge tube (Tube 2) and sonicated at 40°C for 10 minutes. A fluorescent label (AlexaFluor 647 c2-maleimide; ThermoFisher) was then added to Tube 2 at a final concentration of 0.02 mM and vortexed. A photoinitiator (lithium phenyl-2,4,6-trimethylbenzoylphosphinate (LAP); Sigma-Aldrich) was added to Tube 1 at a final concentration of 4.5 mM and vortexed. The entire contents of Tube 2 were then added to Tube 1 and vortexed to form the aqueous component of the emulsion polymerization. The entire 1.5 mL aqueous solution was then emulsified in 5 mL of degassed light mineral oil (Fisher Scientific) with 0.5 wt% Span80 (Sigma-Aldrich) by stirring in a Sigmacote (Sigma-Aldrich) coated glass scintillation vial at 300 rpm. The emulsification was stirred for 30 seconds and then the suspension was exposed to 365 nm UV light at 60 mW/cm^2^ (Omnicure, Lumen Dynamics) for 5 minutes. Stir speed and duration were experimentally determined to yield microspheres with a median diameter of approximately 200 µm^30^.

The reacted product was collected in a 50 mL conical tube, washed with fresh mineral oil, and centrifuged (Thermo Scientific Sorvall ST 40R) at 300 rpm for 3 minutes. This washing step was repeated twice for a total of three mineral oil washes. Following the final mineral oil wash, the hydrogel microspheres were washed once in 100% isopropyl alcohol (Fisher Scientific) and centrifuged at 600 rpm for 5 minutes. The supernatant was discarded and the hydrogel microspheres were stored overnight in sterile phosphate buffered saline (PBS; Cytiva) at 4°C. Microspheres were filtered successively through 200 µm, 100 µm, and 85 µm cell strainers (pluriSelect ÜberStrainer). The microspheres collected in the 85 µm strainer were used for experiments.

#### Isolation of Primary Murine ATII Cells and Fibroblasts

All animal procedures were performed in an AAALAC-accredited facility in accordance with the Guide for the Care and Use of Laboratory Animals^37^. All protocols were approved by the University of Colorado Denver Institutional Animal Care and Use Committee. Male and Female, 8- to 12-week-old, dual-transgenic reporter C57BL/6J mice were bred for use in these experiments. Wildtype littermates were used for alveolar epithelial type II (ATII) cell isolation and viability experiments. Primary murine ATII cells and fibroblasts were isolated from healthy mice by magnetic column isolation as follows.

The lungs of freshly sacrificed mice were perfused intratracheally with a room temperature solution of dispase (5 U/mL; Gibco) and collagenase type I (2 mg/mL; Gibco) in PBS. A 1% low-melting point agarose solution (ThermoFisher UltraPure) was subsequently injected intratracheally and the lungs were covered in ice to form a solidified agarose plug. The lungs were removed and kept on ice in PBS until being transferred to a gentleMACS C Tube (Miltenyi Biotec) with fresh dispase/collagenase solution and incubated on a rotator at 37°C for 20 minutes. The lungs were then mechanically dissociated on a GentleMACS Dissociator (Miltenyi Biotec) using the “m_lung_02.01” C Tube setting and strained to create a single-cell suspension. The cells were counted on a hemacytometer (Hausser Scientific) as the suspension was centrifuged at 1200 rpm for 5 minutes at 4°C. The supernatant was discarded and the pellet was resuspended in a buffer consisting of 0.5% bovine serum albumin (BSA; Sigma-Aldrich) and 2 mM ethylenediaminetetraaetic acid (EDTA; ThermoFisher) in PBS (PEB buffer) at a concentration of 80 µL per 10^7^ total cells.

Endothelial (CD31^+^) and hematopoietic (CD45^+^) cells were removed from the suspension. These cells were magnetically labelled by adding 10 µL CD31 and CD45 antibody-conjugated microbeads (Miltenyi Biotec) per 10^7^ total cells and incubating at 4°C for 15 minutes. Cells were washed in 1 mL PEB buffer per 10^7^ cells, centrifuged at 1200 rpm for 5 minutes at 4°C, and resuspended in 500 µL PEB buffer per 10^8^ cells. The cells were then pipetted onto prepared LS columns (Miltenyi Biotec) fixed in a QuadroMACS separator (Miltenyi Biotec) at 500 µL/column. After the cell suspension had flowed through the column, the columns were washed twice with 3 mL PEB buffer. The CD31^−^/CD45^−^ flowthrough was collected, counted, and resuspended in 90 µL PEB buffer per 10^7^ total cells.

Epithelial cells expressing epithelial cell adhesion molecule (EpCAM^+^) cells were sorted and reserved for experiments. These cells were labelled and isolated in the same way using microbeads conjugated with an EpCAM antibody. EpCAM^+^ cells were removed from the LS column by removing the column from the QuadroMACS separator, placing the column over a 15 mL conical tube, and quickly flowing 5 mL PEB buffer through the column twice. The isolated EpCAM^+^ cells were then counted, centrifuged at 1200 rpm for 5 minutes at 4°C, and resuspended in complete medium (DME/F-12; Cytiva) with 100 U/mL penicillin, 100 mg/mL streptomycin, and 2.5 mg/mL amphotericin B (Life Technologies), supplemented with 10% fetal bovine serum (FBS; ThermoFisher).

Primary murine fibroblasts were isolated from dual-transgenic reporter C57BL/6J mice following the same protocol. After the removal of CD31^+^/CD45^+^ cells, alveolar fibroblasts expressing platelet derived growth factor receptor alpha (PDGFRα^+^) were labelled and isolated as described above ^38^. The isolated PDGFRα^+^ cells were then counted, centrifuged at 1200 rpm for 5 minutes at 4°C, and resuspended in complete growth media.

#### Formation of 3D Acinar Structures

The isolated ATII cells were collected, counted, and magnetically labeled to enable magnetic aggregation in 3D acinar structures. Cells were resuspended in complete media at 2.5×10^6^ cells/mL and NanoShuttle (Greiner Bio-One) was added at a concentration of 250 µL/mL media. The mixture was mixed by pipetting up and down and centrifuged at 1200 rpm at 4°C for 5 minutes. The mixture was mixed and centrifuged two more times for a total of three centrifugations. The magnetically labeled ATII cells were mixed with microspheres at 250 cells/microsphere and aliquoted into an ultra-low adhesion 24-well plate at 250 microspheres/well. Complete media supplemented with 10% FBS was added to each well up to 350 uL total volume and a 24-Well Levitating Drive (Greiner Bio-One) was placed over the plate. The plate was then incubated at 37°C for 72 hours on an orbital shaker plate to allow aggregation of the 3D acinar structures.

#### Preparation of Embedding Hydrogels

The 3D acinar structures were embedded in either a soft or stiff PEG-NB hydrogel to represent healthy or diseased lung tissue. The embedding hydrogel contained a 10 kg/mol PEG-NB backbone and an MMP2-degradable dithiol crosslinker (KCGPQGIWGQCK; Genscript) to enable remodeling by fibroblasts^39^. The weight percentages and crosslinker ratios of the embedding gels were determined experimentally through parallel plate rheology to mimic the elastic modulus (E) of healthy (5.0 wt% PEG-NB, r = 0.7, E = 2.7 kPa) and fibrotic (7.5 wt% PEG-NB, r = 0.8, E = 18.1 kPa) lung tissue (N = 8). Embedding hydrogels were prepared by dissolving the 10 kg/mol PEG-NB backbone, MMP2-degradable crosslinker, and biologically active peptides (CGRGDS and CGYIGSR) individually in complete media supplemented with 10% FBS and sonicating for 10 minutes at 40°C. The PEG-NB was combined at the appropriate final weight percentage with the biologically active peptides, both at a 2 mM final concentration, and vortexed (Tube 1). The crosslinker was combined at the appropriate r-ratio with 15 mM TCEP and sonicated for 10 minutes at 40°C (Tube 2). LAP was added to Tube 1 at a final concentration of 4.5 mM and vortexed, and the entire contents of Tube 2 were added to Tube 1. The pH of this prepared stock solution was then measured using color bonded pH test papers (Cytiva) and neutralized with 3 M potassium hydroxide (Sigma-Aldrich) and 1 M hydrochloric acid. Freshly isolated primary murine fibroblasts were added to complete media supplemented with 10% FBS. This cell suspension was then added to the hydrogel stock solution for a final concentration of 60,000 cells per sample (6,000 cells/µL stock solution) to bring the stock solution up to the desired final gel volume.

#### Assembly of 3D Lung Models

The magnetic levitating drive was removed and the 24-well plate containing the microspheres and ATII cells was placed on a 24-Well Holding Drive (Greiner Bio-One). The media was manually aspirated from each well and the 3D acinar structures were circled with a hydrophobic pen. 10 µL of the prepared embedding hydrogel and fibroblast solution was pipetted onto each structure using a positive displacement pipette. The samples were exposed to 365 nm UV light at 10 mW/cm^2^ (Omnicure, Lumen Dynamics) for 5 minutes. The 3D lung models were then transferred to a new 24-well plate with complete media supplemented with 10% FBS and incubated at 37°C. Samples embedded in soft embedding hydrogel were kept in Costar 24-well transwell inserts (Corning) to minimize exposure to mechanical forces during media changes.

### Characterization of 3D Lung Models

#### Hydrogel Characterization

Evaluation of hydrogel mechanical properties was performed by parallel plate rheology. Briefly, hydrogel samples (height = 1 mm; diameter = 8 mm) were prepared according to the procedure described above and swelled overnight in PBS at room temperature. The samples were then trimmed as necessary and fitted onto a Discovery HR2 rheometer (TA Instruments) between an 8 mm diameter parallel plate geometry and a Peltier plate set to 37°C. The geometry was lowered until 0.03 N axial force was applied. The gap distance was noted and decreased until the storage modulus measurement (G’) plateaued. The percent compression at plateau of the specific hydrogel was used for each subsequent measurement for that sample type^40^. All samples were subjected to frequency oscillatory strain with a range of 0.1 to 100 rad/s at 1% strain. The elastic modulus (E) was calculated using rubber elasticity theory, assuming a Poisson’s ratio of 0.5 for bulk measurements of the elastic hydrogel polymer network^41^.

#### Characterization of Hydrogel Microspheres

Hydrogel microsphere emulsion polymerization conditions were determined experimentally to optimize microsphere diameter. Multiple stir speeds were tested across multiple emulsion times prior to UV exposure and the size distribution of the resulting microspheres was analyzed on an upright epi-fluorescent microscope (BX-63; Olympus). Microsphere diameters were measured using ImageJ and a histogram was generated from each experimental condition. Individual microspheres were identified on ImageJ using the watershed function and subsequently analyzed. The resulting major axis measurement of each microsphere was taken to be the diameter. The condition resulting in spheres with a median diameter closest to 200 µm (300 rpm, 30 seconds; median diameter = 171 ± 42 µm) was chosen for this experiment.

#### Cellular Viability

Long-term viability of the primary murine ATII cells and fibroblasts co-cultured within both the healthy and fibrotic 3D lung models was confirmed with a live/dead staining assay (Millipore Sigma QIA76). 3D lung models were rinsed once with PBS and then incubated in a 1:1000 dilution each of Cyto-dye (green) and propidium iodide (red) for 40 minutes at 37°C. Samples were rinsed in PBS and then placed in a solution of 10% FBS in PBS for immediate imaging on a 3i MARIANAS inverted spinning disk confocal microscope. Three 50 μm z-stacks were acquired per sample, with three samples per condition. Positive cells in each channel were quantified using Fiji (ImageJ) by producing a maximum projection of the z-stack, thresholding, and counting particles. Data are presented as N = 3 where each sample is the average of three images.

#### Visualization of Cellular Distribution in 3D Lung Models

Whole 3D lung models were rinsed in PBS, fixed for 30 minutes in 4% paraformaldehyde (PFA; Electron Microscopy Sciences) in PBS, and permeabilized for 30 minutes in 0.5% TritonX-100 in PBS (Fisher). The samples were then blocked for 1 hour in 3% BSA in PBS before overnight incubation at 4°C with rabbit anti-SFTPC (ThermoFisher PA5-71680, 1:25), hamster anti-Podoplanin (ThermoFisher MA5-18054, 1:100), and rat anti-Cytokeratin 8 (Developmental Studies Hybridoma Bank TROMA-I, 1:25). Samples were washed three times with 0.1% Tween-20 (Fisher) in PBS, incubated for 1 hour at room temperature with AlexaFluor 488 goat anti-rat, AlexaFluor 647 goat anti-hamster, and AlexaFluor 555 goat anti-rabbit (ThermoFisher, 1:400), then again washed three times with 0.1% Tween-20 in PBS. Samples were stained with 5 μg/mL Hoechst (Tocris) in PBS for 30 minutes, washed three times with PBS, and then imaged on an upright, epifluorescent microscope (Olympus, BX-63). For each sample, three 60 µm z-stacks were acquired with the 10x objective. Image processing was performed by first running a Weiner deconvolution and generating a maximum projection before thresholding and counting particles in Fiji (ImageJ).

#### Cellular Activation

3D lung models were generated using fibroblasts isolated from dual-reporter mice. These cells express GFP under the control of the collagen 1 alpha chain 1 (Col1a1) promoter promoter and RFP under the control of the alpha smooth muscle actin (αSMA) promoter, allowing for visualization of myofibroblast activation using endogenous labels. Fibroblasts were cultured either alone or in co-culture with ATII cells and allowed to recover in complete media with 10% FBS for one week, after which they were cultured in low-serum activation media (complete media with 1% FBS) for an additional 1-2 weeks. At each time point, six samples per condition (soft and stiff embedding hydrogel, with and without ATII cells) were processed for imaging analysis. Whole samples were incubated with 5 μg/mL Hoechst in PBS for 30 minutes at 37°C, washed once with PBS, and then kept in complete media during imaging on a 3i MARIANAS inverted spinning disk confocal microscope. Three 100 μm z-stacks were acquired per sample and were quantified in Fiji by producing a maximum projection of each channel of the z-stack, thresholding, and counting particles. Data are presented as N = 6 where each sample is the average of three images.

#### RNA Isolation and qPCR Array

RNA was isolated from the 3D lung models after 21 days of incubation. Three samples were pooled in RNAse-free 1.5 mL microcentrifuge tubes and rinsed in sterile PBS for 5 minutes on a rocker. The PBS was then removed and the samples were flash-frozen by submerging the tubes in liquid nitrogen. Samples were manually homogenized with an RNAse-free plastic pestle and by pipetting during the thawing process. Samples were then incubated in 500 µL serum-free media with 2 mg/mL collagenase at 37°C for 10 minutes, or until the embedding hydrogels had visibly degraded. Next, 500 µL TRIzol reagent (Life Technologies) was added to each tube and the samples were mixed by pipetting and incubated on ice for 5 minutes. 100µL 1–bromo–3–chloropropane (BCP; Fisher Scientific) was then added to each sample. The samples were vortexed, incubated at room temperature for 10 minutes, then cooled on ice and centrifuged at 14,000 g for 15 minutes. The clear layer of each sample was transferred to a new RNAse-free 1.5 mL microcentrifuge tube with 500 uL of isopropyl alcohol, vortexed, and incubated at room temperature for 10 minutes. The samples were cooled on ice and centrifuged at 14,000 g for 10 minutes. The supernatant was discarded and 1 mL 70% ethanol (Decon Laboratories) was added. The samples were vortexed and centrifuged at 14,000 g for 5 minutes. The supernatant was discarded and the samples were allowed to air dry. The dried pellets were resuspended in 16 µL of RNAse-free water (Qiagen). RNA quantity and purity were measured on a BioTek plate reader using a Take3 Micro-Volume Plate. Purity was assessed based on the ratio of absorbance readings taken at 260 nm and 280 nm (A_260_/A_280_).

The isolated RNA was then analyzed using a mouse fibrosis specific RT^2^ Profiler PCR Array (Qiagen), which measures the expression of 84 genes known to be involved in the progression of fibrosis. 40 U RNAse inhibitor (Sigma-Aldrich) was added to the eluted RNA. cDNA was created from 3.6 µg RNA using a Qiagen RT^2^ First Strand Kit.

#### Statistical Analysis

Viability experiments comparing soft versus stiff embedding hydrogel were assessed by two-tailed t-test at each timepoint. Activation experiments were analyzed by two-way analysis of variance (ANOVA) with Tukey’s Test for multiple comparisons. Gene expression analysis was performed on the RT^2^ Profiler PCR Array Data Analysis Webportal (Qiagen). Normalized gene expression (2^(-ΔCT)) was calculated relative to the housekeeping gene B2M and genes with a fold regulation of at least two in any experimental group relative to the control were incorporated into the heatmap (Figure 6).

## Results & Discussion

### 3D lung models recapitulate pulmonary micro-architecture and elastic modulus

We present a strategy for engineering 3D lung models using PEG-NB hydrogels that enables researchers to study crosstalk between epithelial cells and fibroblasts as well as cellular responses to changes in mechanical properties within precisely defined 3D geometries that recapitulate lung tissue. First, hydrogel microspheres were generated to serve as a template for arranging epithelial cells into geometries that mimic lung micro-architecture. PEG-NB macromers (40 kg/mol) were combined with an MMP3-degradable crosslinker, thiolated magnetic nanoparticles (MNPs) to enable magnetic aggregation into acinar structures, and peptide sequences from fibronectin (CGRGDS) and laminin (CGYIGSR) to promote cell attachment and survival, in a photoinitiated emulsion polymerization procedure (Figure 1a). The same thiol-ene click chemistry incorporated a fluorescent label to allow visualization and measurement of the microspheres (Figure 1b). Following emulsion polymerization, the resulting hydrogel microspheres were filtered with 100µm and 85µm filters to select a population that displayed sizes matching the average diameter of the human alveolus (median diameter = 171 ± 42.1 µm; Figure 1c)^24^. Parallel plate rheology confirmed that the elastic modulus of these hydrogel microspheres (4.62 ± 1.01 kPa; Figure 1d) matched values reported for healthy lung tissue (2-5 kPa)^22^.

**Figure 1.**
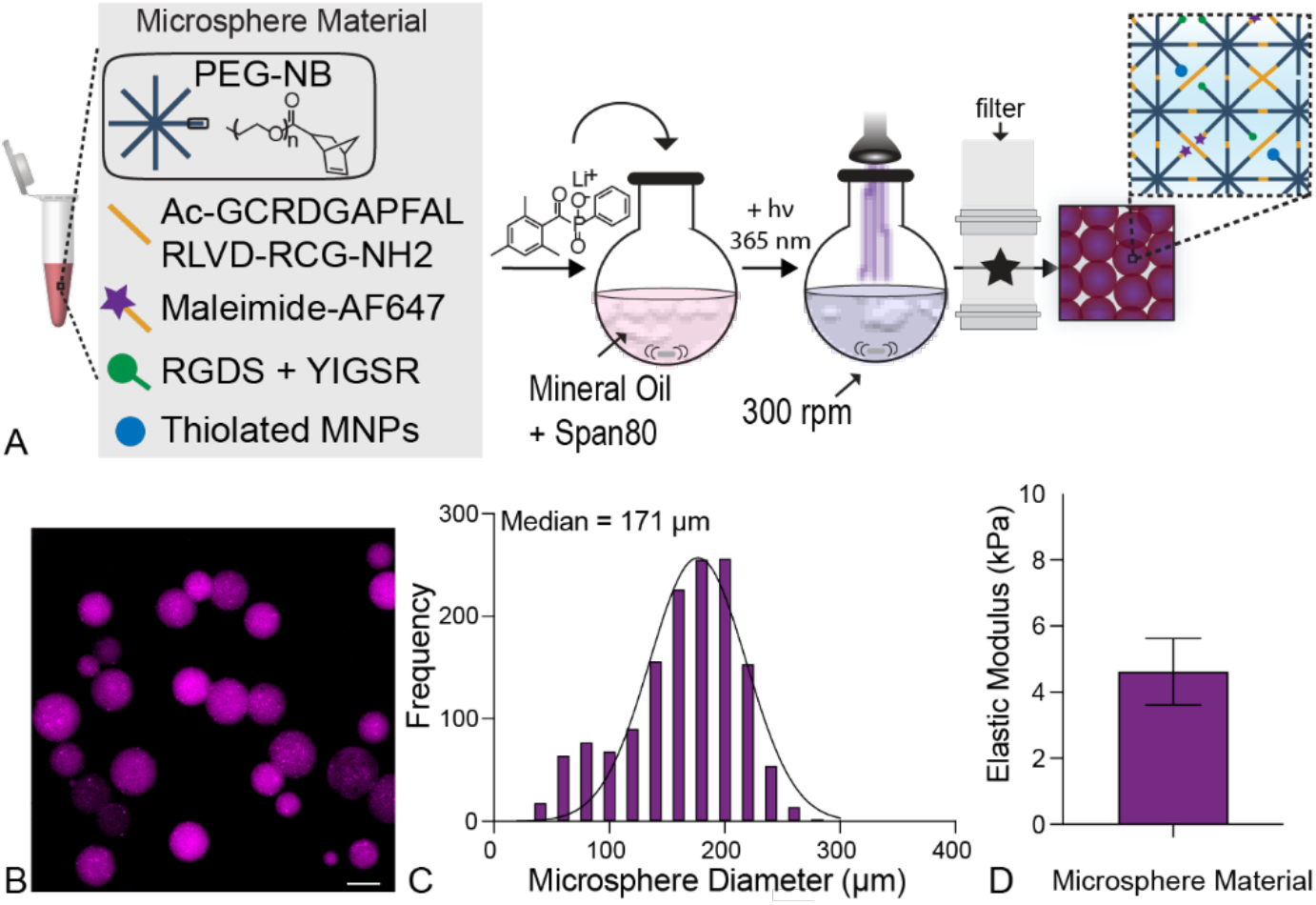
Emulsion polymerization produced hydrogel microspheres for templating alveolar micro-architecture. A) An overview of the microsphere fabrication process detailing chemical components and emulsion parameters. Created with BioRender.com. B) Representative fluorescent image of polymerized microspheres showed spherical geometry. Scale bar, 100 µm. C) Image analysis of microsphere diameter from fluorescent microscope images demonstrated that collecting 85-100 µm filtrate yielded physiologically relevant microspheres sizes (median diameter = 171 ± 42 µm; N = 4). D) Parallel plate rheology confirmed that microsphere stiffness was within the range of healthy lung tissue (E = 4.62 ± 1.01 kPa; N = 8).

To form higher-order acinar structures, hydrogel microspheres were coated with primary murine ATII cells and aggregated under a magnetic field for three days, resulting in even distribution of ATII cells around tightly clustered microspheres (Figure 2a). These 3D acinar structures were then embedded in another PEG-NB hydrogel laden with primary murine fibroblasts to create 3D lung models (Figure 2a). The embedding hydrogel was comprised of the PEG-NB macromer crosslinked with an MMP2-degradable peptide sequence, along with the same fibronectin and laminin-derived adhesion peptides. The peptide crosslinker was designed to allow environmental remodeling by MMP2-expressing fibroblasts. The ability of fibroblasts to interact with the extracellular microenvironment and spread is critical for activation to the myofibroblast phenotype within fibrous synthetic composite hydrogels^42^ (Figure 2b). The formulation of the embedding hydrogel was tailored to produce either a soft embedding hydrogel (E = 2.7 ± 0.31 kPa) to mimic healthy lung (E = 2-5 kPa ^22^), or a stiff embedding hydrogel (E = 18.1 ± 1.34 kPa) to mimic fibrotic lung (E = 10-30 kPa^22^; Figure 2c) as measured by parallel plate rheology. The result of this microsphere aggregation and embedding procedure was the production of a 3D lung model in which primary murine ATII cells formed alveoli-like structures while surrounded by an environment that reproduced the mechanical microenvironment and cellular landscape of pulmonary interstitial tissue.

**Figure 2.**
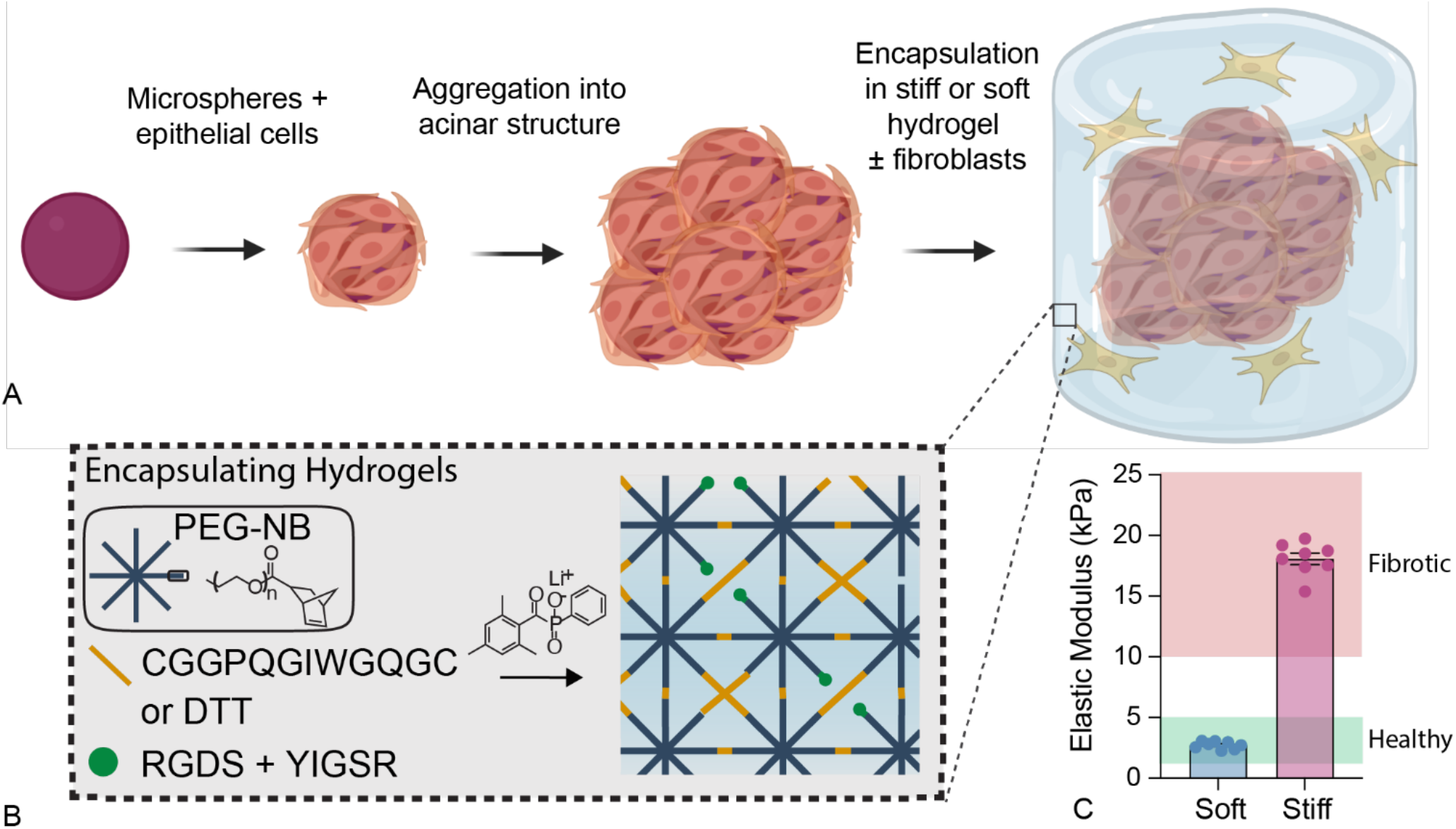
Formation of 3D acinar structures. A) Overview of the 3D acinar structure fabrication process. Created with BioRender.com. B) Schematic depicting the embedding hydrogel formulation and polymerization process. These materials were crosslinked with an MMP2-degradable dithiol crosslinker to enable fibroblast-mediated matrix remodeling. C) Parallel plate rheology confirmed that embedding hydrogel formulations recapitulated the elastic modulus of healthy (E = 2.7 ± 0.31 kPa) or fibrotic lung tissue (E = 18.1 ± 1.34 kPa). N = 8.

Previous studies have reported culturing lung cells on hydrogel microspheres to model alveolar micro-architecture. Lewis *et al*. fabricated photodegradable PEG-based hydrogel microspheres (mean diameter = 120 ± 70 µm) and coated these templates with epithelial cells. Single microspheres were embedded in a stiff hydrogel (E ∼ 20kPa) and exposure to ultraviolet light degraded away the microsphere templates, creating a cyst that mimicked the structure of a single alveolus^24^. By encapsulating a fibroblast cell line in the embedding hydrogel, they subsequently demonstrated that A549 cancer cells could activate fibroblasts in this 3D model. Our model relies on similar PEG-based microspheres, but aggregated into larger acinar structures and with the incorporation of primary lung cells, rather than cell lines. Similarly, Wilkinson *et al*. used a custom bioreactor to aggregate human fetal lung fibroblasts around collagen-functionalized alginate microspheres (diameter = 161 ± 80 µm) to create mesenchymal lung organoids that formed an extended tissue network more fully reproducing 3D lung micro-architecture and allowing for the study of transforming growth factor-β (TGF-β) mediated fibroblast activation in the presence of additional cell types^26^. In contrast to the static natural materials used to form these hydrogel microspheres, our PEG-based hydrogels are highly customizable, allowing us to also assess cellular responses to changing environmental mechanics. In this way, we combined the tunability of PEG hydrogels for microsphere fabrication with a magnetic aggregation step to create a novel 3D model of higher-order lung structure. This model contains primary epithelial cells that can be embedded with fibroblasts inside hydrogels to mimic either healthy or diseased lung parenchyma.

### Cells maintain viability and undergo differentiation within 3D lung models

To assess the long-term viability of cells within 3D lung models, constructs were maintained in culture for up to three weeks and the percent of live cells was measured weekly via live/dead nuclear staining. The day after embedding (week 0), aggregate viability was initially lower than subsequent days and displayed high variability (71.1 ± 18.5% for soft and 51.6 ± 14.4% for stiff). After one week in culture the cells recovered and then maintained high viability between 77.4% and 90.7% for both embedding hydrogel conditions out to three weeks (Figure 3a). Critically, cells in the center of aggregates maintained high viability across all timepoints, indicating that nutrient and gas diffusion proceeds freely throughout the 3D structure (Figure 3b).

**Figure 3.**
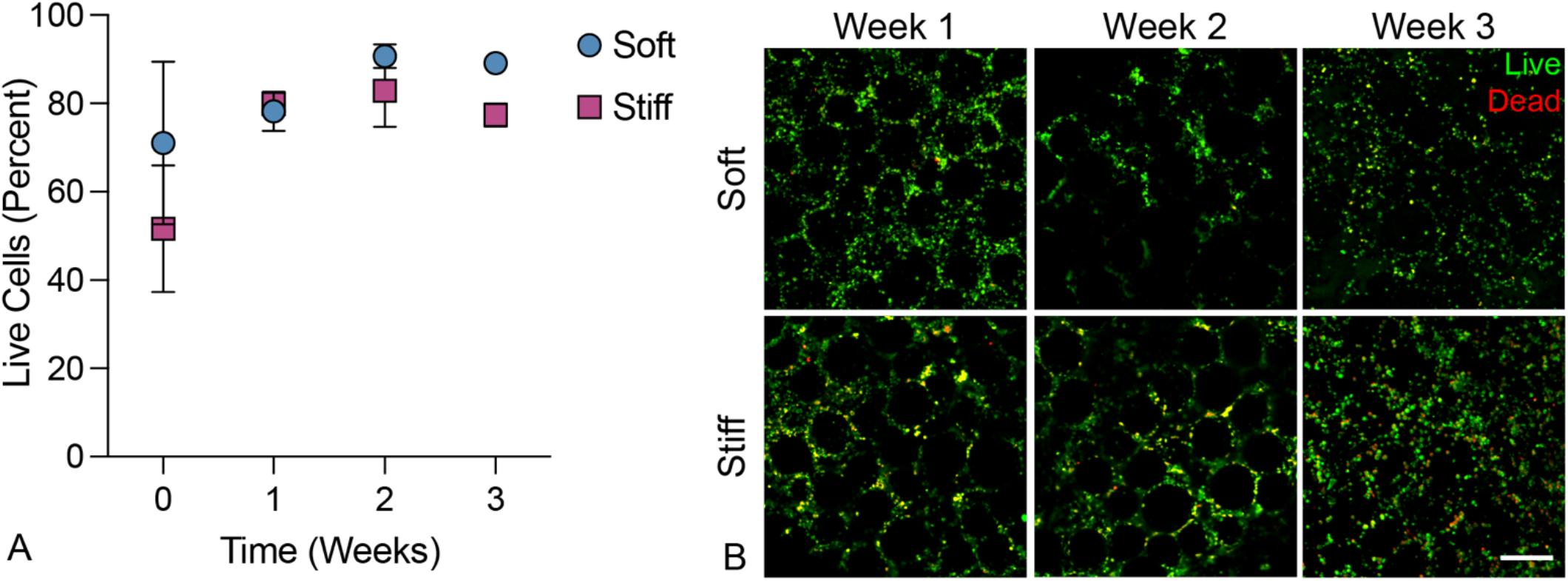
3D Lung models supported cell viability over three weeks in culture. A) 3D lung models supported cell viability out to at least 3 weeks post-embedding in both hydrogel conditions. B) Representative confocal images (50 µm z-stacks displayed as a maximum intensity projection) showed even distribution of live cells throughout 3D lung models and preserved pulmonary architecture. Scale bar, 200 µm. N = 3.

*In situ*, ATII cells possess the ability to differentiate into ATI cells, changing their morphology, function, and gene expression in response to epithelial injury. This natural tendency for ATII cells to drift toward an ATI phenotype has also been observed *in vitro* in 3D hydrogel models^24^. It is well documented that the lung epithelium is heavily involved in the regulation of pulmonary fibrosis. Healthy epithelium can secrete antifibrotic factors such as prostaglandin-E2 (PGE2) and bone morphogenetic protein-4 (BMP4) that restrain fibroblast activation and help maintain healthy lung morphology^21, 43^. Conversely, damaged epithelium secretes profibrotic factors, including TGF-β, PDGF, and CTGF, that induce myofibroblast activation and fibrotic progression^6, 7^. Epithelial damage also induces differentiation of ATII cells into ATI cells as part of normal, healthy re-epithelialization. During fibrosis, a population of transitional ATII cells persists and contributes to a positive feedback loop of fibrotic progression^5^. We assessed epithelial cell identity within aggregates over time by immunofluorescence staining for an ATII cell marker, surfactant protein C (SFTPC), a transitional ATII-ATI marker, keratin-8 (KRT8), and an ATI cell marker, podoplanin (PDPN; Figure 4a). While the majority of seeded epithelial cells (approximately 85%) initially expressed SFTPC, over the course of three weeks a population of PDPN+ ATI cells emerged, indicating differentiation over time. Interestingly, at the initial time point, 10-15% of seeded cells were already expressing the transitional marker KRT8, and this population expanded up to 20-25% by week 3 (Figure 4b). These shifts in epithelial cell population mimic the sustained transitional cell phenotype observed in pulmonary fibrosis, where transitional ATII cells are known to emerge in response to epithelial injury and share pro-fibrotic hallmarks with various other pathological states^44^. These transitional cells have been found to display phenotypes similar to epithelial-mesenchymal transition^45^ and cellular senescence^46^, particularly in terms of increased TGF-β signaling. The primary murine ATII cells included in this model started to undergo this transition, resulting in a mixed population of ATII and ATI cells alongside cells that are arrested in this transitional pathologic state. Critically, these aggregates all contained PDGFRα-expressing fibroblasts, an alveolar fibroblast population that has been noted to promote both ATII cell self-renewal and differentiation to ATI cells^38^. While the transitional epithelial cell phenotype has, to date, largely been observed *in vivo*, emerging *in vitro* data suggests that fibroblasts also play a critical role in initiating and maintaining this unique epithelial cell state^47^. Taken together, these data demonstrate the capacity of this *in vitro* co-culture model to recapitulate the cellular heterogeneity that is a hallmark of fibrosis *in vivo*.

**Figure 4.**
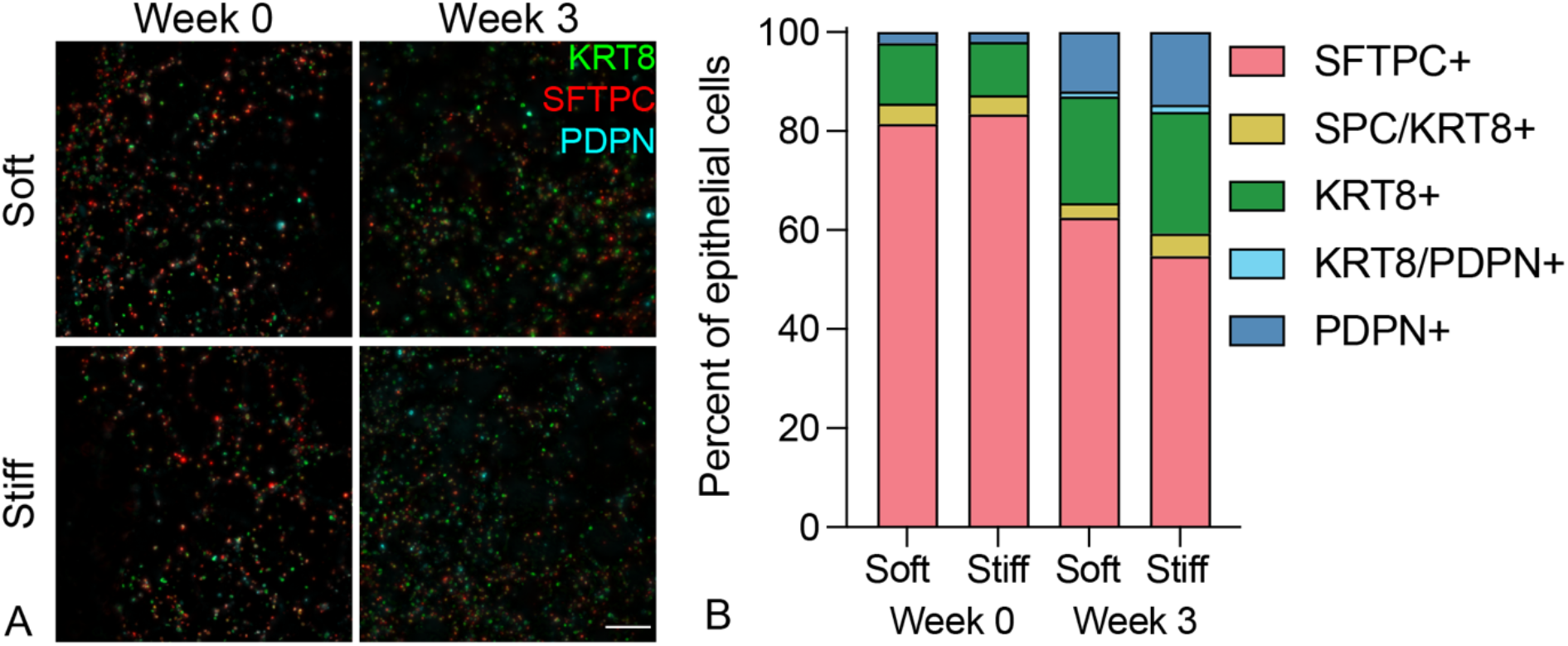
Epithelial cell identity shifted over time. A) Representative fluorescent images of immunostaining of ATII (SFTPC), ATI (PDPN), and transitional (KRT8) markers. B) Image analysis showed the majority of the ATII cell population retained that identity, with some ATII cells transitioning to an ATI or ATII-ATI transitional phenotype at 3 weeks post-embedding (N = 3). Scale bar, 100 µm.

### Presence of ATII cells promotes fibroblast activation in stiff environment

Fibroblasts are the central drivers of fibrotic progression because these cells synthesize and remodel fibrotic ECM in a positive feedback loop of cellular activation and tissue stiffening^48^. The importance of epithelial-mesenchymal crosstalk to this fibrotic activation has been observed *in vivo* but is still lacking within *in vitro* models. To interrogate the effects of both microenvironmental stiffness and epithelial cell interactions on fibroblast activation in the context of IPF, fibroblasts were isolated from dual-transgenic reporter mice. These reporter cells express red fluorescent protein (RFP) under the control of the alpha smooth muscle actin (αSMA) promoter and green fluorescent protein (GFP) under the control of the collagen 1 alpha chain 1 (Col1a1) promoter, offering a visual indication of cellular activation and transition toward the active myofibroblast phenotype^49^. Fibroblasts were cultured in four conditions: around 3D acinar structures, either with or without epithelial cells incorporated, and within either soft or stiff embedding hydrogels, with activation assessed at weeks 0, 2, and 3 in culture. Fibroblasts cultured with epithelial cells in a stiff extracellular environment showed significantly higher expression of αSMA and Col1a1 after 3 weeks compared to co-culture in a soft extracellular environment or fibroblasts in monoculture (Figure 5a). In particular, the presence of epithelial cells caused an early expansion of αSMA+ myofibroblasts by week 2 (Figure 5b). By week 3, the dual stimulus of epithelial cell presence and microenvironmental stiffness dramatically increased the population of activated fibroblasts (Figure 5c). These results supported the hypothesis that the combination of mechanical and cellular cues is critical to promoting a fibrotic response, more so than either mechanical or cellular cues alone.

**Figure 5.**
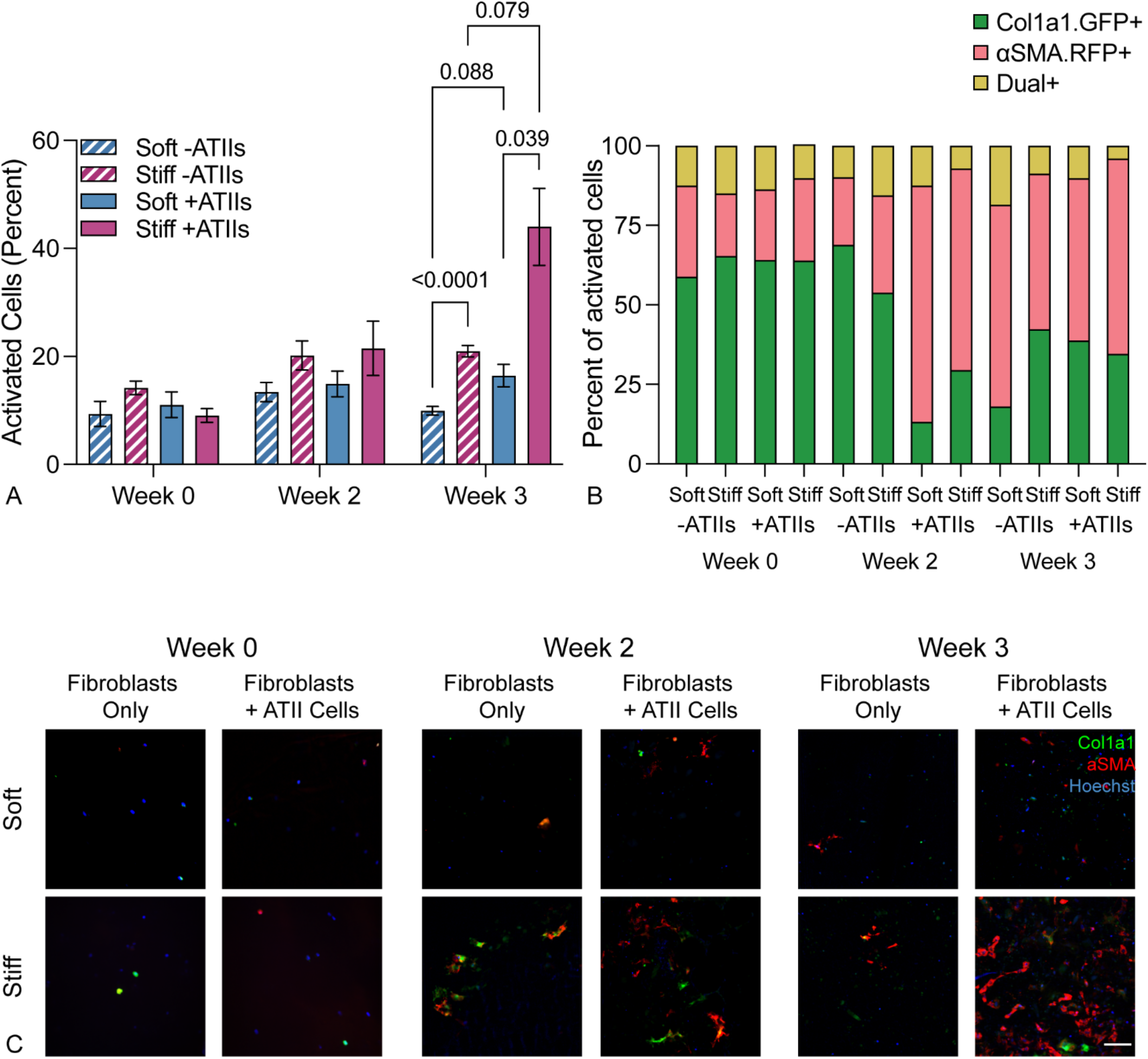
Fibroblasts were activated in response to stiffness and epithelial cell presence. A) Fibroblast activation assessed up 3 weeks post-embedding revealed that fibroblast-epithelial cell co-culture and a stiff extracellular environment synergistically promoted fibroblast activation as measured by the percentage of αSMA and/or Col1a1 expressing reporter fibroblasts in culture (N = 6). B) Within the activated cell population, the proportion of αSMA+ myofibroblasts increased over time in all conditions, but increased more quickly in co-culture conditions, suggesting that the presence of epithelial cells provokes a more rapid fibrotic response than mechanical cues alone (N = 6). C) Representative confocal images of dual reporter fibroblasts within 3D lung models showed increased expression of activation reporters in co-culture within a stiff extracellular environment, particularly at three weeks. Scale bar, 100 µm.

Prior studies have demonstrated fibroblast activation in response to mechanical cues. Fibroblasts cultured on a 2D hydrogel with an established stiffness gradient exhibited increased spreading, proliferation, and migration as the elastic modulus of the substratum increased^50^. The relationship between environmental stiffness and fibroblast activation is more complex in 3D models, where factors such as the density or deformability of the surrounding environment also play a role^42^. In our 3D hydrogel model, we demonstrate that increased microenvironmental stiffness alone can increase fibroblast activation, but that epithelial cells also play a synergistic role in this activation (Figure 5a). Cellular crosstalk between fibroblasts and epithelial cells has been studied in numerous ways. Studies that have used conditioned media^44^ or trans-well co-culture^51^ to demonstrate the reciprocal crosstalk between damaged epithelium and activated fibroblasts have implicated key signaling pathways in the development of IPF, but still rely on 2D cell culture, often on substrates with non-physiologically mechanical properties, resulting in abnormally high fibroblast activation. In an organoid co-culture model, healthy epithelium inhibited the activation of fibroblasts through BMP signaling, even in the face of fibrotic stimulation with exogenous TGF-β. Interestingly, this mitigated activation in the presence of TGF-β was only observed in 3D culture, not 2D. These data support the concept that lung epithelium can play a critical regulatory role in the development of fibrosis and also highlight the power of studying cellular behavior in geometrically relevant 3D culture systems^21^. Here, fibroblasts and epithelial cells were co-cultured in a micropatterned 3D environment and the presence of epithelial cells enhanced fibroblast activation. Critically, these results, along with immunostaining, suggested that the epithelial cells are not recapitulating healthy epithelium, but are reproducing aspects of fibrotic epithelium, with an injury response resulting in differentiation of ATII cells to ATI cells and the persistence of a transitional cell population (Figure 4b). This mixed epithelial cell population, in concert with increased microenvironmental stiffness, results in a robust model of pathological fibroblast activation, highlighting the importance of models that allow researchers to control and study both cell-matrix and cell-cell interactions.

### Stiff PEG-NB microenvironment co-culture promotes upregulation of profibrotic genes

qRT-PCR analysis of 84 genes known to be involved in the progression of fibrosis was performed to further assess fibroblast activation and the fibrotic phenotype of cells within 3D lung models featuring different mechanical properties. Results revealed differences in the expression of several gene networks in response to environmental stiffness and the presence of epithelial cells (Figure 6). In fibroblast-only cultures, a stiff microenvironment induced upregulation of genes involved in ECM remodeling and integrin signaling, suggesting that the cells sense this external stimulus and respond by expressing factors that would facilitate interactions with the extracellular environment. Interestingly, prior studies in 3D cultures of fibroblasts aimed at the study of IPF demonstrated that interactions with the extracellular microenvironment were critical to the activation of myofibroblasts. Matera *et al*. showed that fibroblasts encapsulated within a cell-degradable hydrogel acquired a myofibroblast phenotype in response to TGF-β stimulation, but this effect was abrogated by treatment with an MMP inhibitor, indicating that the ability of fibroblasts to remodel their surrounding environment is critical to their activation^42^. In another model using decellularized slices of lung tissue as a 3D scaffold for cell culture, Booth *et al*. demonstrated that ECM derived from fibrotic lung was sufficient to activate myofibroblasts, even in the absence of overall differences in TGF-β signaling^52^. Both these data and ours highlight the critical role that cell-matrix interactions have in fibroblast activation.

**Figure 6.**
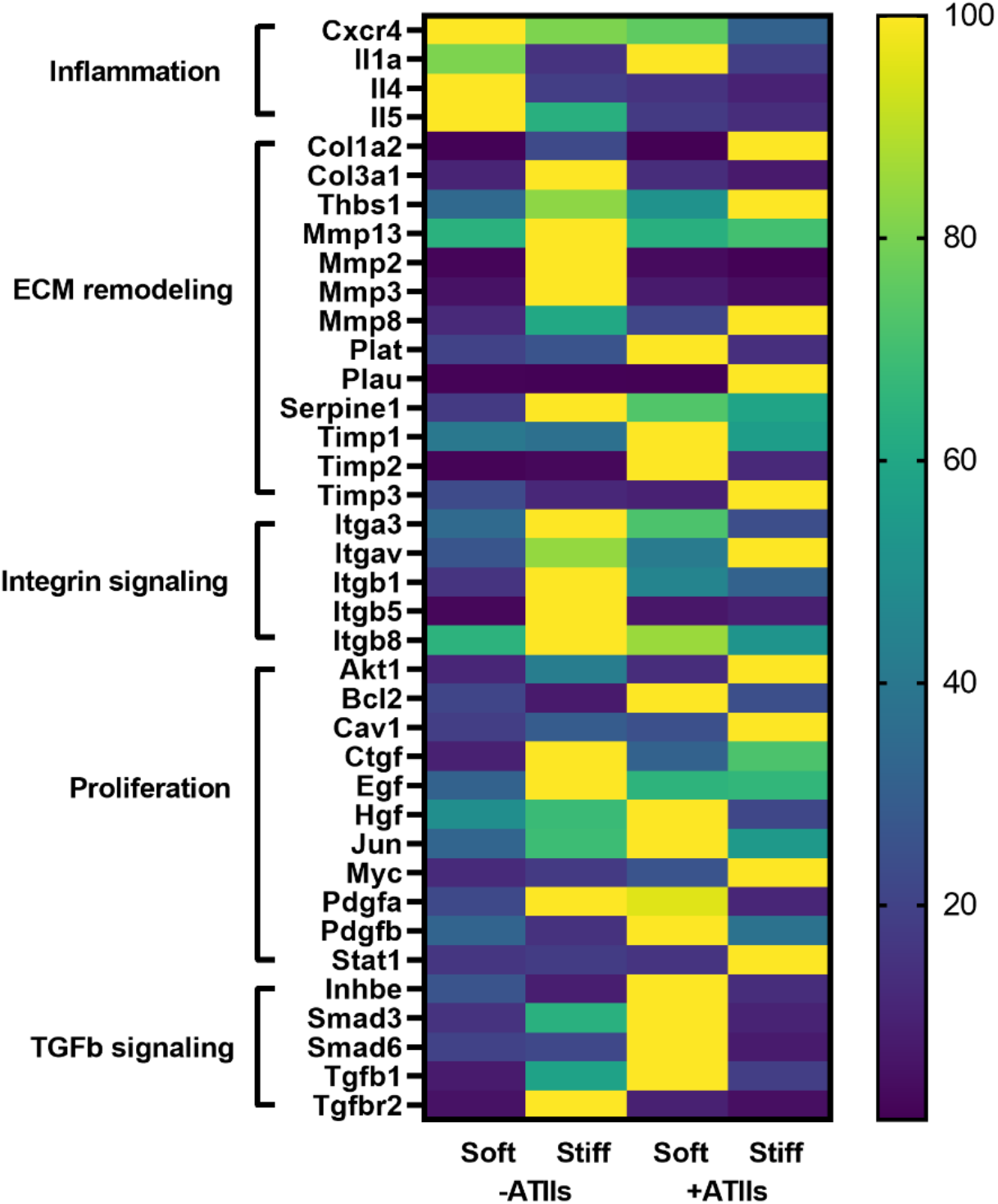
Fibrotic gene network analysis in 3D lung models. Expression of 84 fibrosis-associated genes was assessed using a Qiagen RT^2^ Profiler PCR array. A subset of genes with a fold change greater than two in at least one condition relative to the soft, fibroblast-only condition is displayed. Within each gene, expression relative to the housekeeper (2^(−ΔCT)) was normalized across the four groups as a percent of maximum expression. This analysis revealed coordinated regulation of functionally related genes based on cellular co-culture and microenvironmental stiffness. Specifically, stiffness tended to promote the expression of genes related to cell-matrix interactions, while presence of epithelial cells created a pro-proliferative environment with increased TGF-β signaling (N = 3).

Relative to fibroblast-only culture in soft microenvironments, co-culture with epithelial cells resulted in increased expression of genes related to TGF-β signaling and cellular proliferation. Although this system did not differentiate whether these genes were being produced by fibroblasts or the epithelial cells, it is suggestive of the creation of a pro-proliferative and pro-activation environment as the result of co-culture. Similarly, Suezawa *et. al*. treated organoids containing both stem-cell derived alveolar epithelial cells and primary fibroblasts with bleomycin, resulting in fibrotic gene expression and functional phenotypes that could not be recapitulated in monocultures of either cell types^27^. These results are in agreement with our own, even though our model did not require the addition of an exogenous pro-fibrotic treatment like bleomycin. Interestingly, the combined condition of co-culture in a stiff microenvironment showed a unique pattern of upregulation of genes across these groupings of cell-ECM and cell-cell interactions. Some of these unique changes may have been a result of direct interactions of epithelial cells with the altered environment. Kim *et. al*. demonstrated that biochemical differences in the ECM impacted the differentiation of epithelial cells, in that alveolar epithelial cells provided with exogenous TGF-β and cultured on fibronectin showed acquisition of epithelial-to-mesenchymal transition markers while those cultured on Matrigel, a different combination of matrix proteins, did not^45^. In our model, we provide epithelial cells with different mechanical inputs, which may also alter injury and stress responses^53^. There is also likely a complex interplay between fibroblasts, epithelial cells, and the microenvironment that contributes to the unique gene expression patterns observed in this 3D co-culture model and could not be recapitulated by more reductionist systems.

One evident shortcoming of this reductionist model comes from the fact that most of the conditions tested displayed downregulation of inflammation-related genes, which play a critical role in the maintenance and injury response phenotypes of alveolar epithelial cells^54^. Incorporation of different fibroblast subsets^38^ and/or inflammatory cells into future 3D lung models could better recapitulate this aspect of fibrotic disease. Still, the findings presented here point to the value of this model in studying both cellular crosstalk between pulmonary fibroblasts and epithelial cells, as well as fibrotic matrix remodeling in a 3D culture model that provides a more relevant and impactful platform for the *in vitro* study of fibrotic disease.

## Conclusion

A significant need exists for improved *in vitro* models of IPF that take into account the variety of factors that contribute to fibrogenesis. Here, we demonstrate the fabrication and validation of a novel *in vitro* model of IPF using mechanically tunable hydrogels polymerized in physiologically relevant alveolar geometries. Magnetically aggregating these hydrogel microspheres into a 3D acinar structure created a more geometrically accurate distal lung model than has previously been established. Incorporating a fibroblast-epithelial cell co-culture along with control over extracellular mechanical cues into this 3D lung model enabled the study of multiple profibrotic factors working in concert with one another, creating a more complete *in vitro* distal lung model than is currently available. This model accurately reproduced the geometric and mechanical properties of native lung tissue and supported cell viability out to at least 3 weeks. Interrogation of the epithelial cells in this model found evidence of ATII cell differentiation into ATI cells, with a persistent population of transitional cells, as has been observed in fibrosis *in vivo*. The fibroblasts in this model displayed significant increases in activation and unique changes to gene expression patterns in response to the presence of epithelial cells and mechanical signals from the extracellular environment, suggesting that the combination of cellular interactions and mechanical cues generated a more profound fibrotic response than either factor alone. The initial success of this model also points toward future improvements that could increase its relevance in the study of human disease. In addition to fibroblasts and epithelial cells, we could also incorporate inflammatory and/or endothelial cells in order to recapitulate the facets of IPF mediated by these cell types. There is also considerable potential in the tunability of our hydrogel materials. Future studies could alter the biochemical environment present in this model by incorporating additional peptide sequences or even ECM-derived protein components. Here, we have built and validated these 3D lung models using murine cells and future studies will incorporate human cells to improve translational investigation of IPF signaling and/or responses to putative therapeutics. Overall, these findings demonstrated that hydrogel-based 3D lung models can serve as a powerful *in vitro* tools for integrating the multitude of complex processes that underlie the progression of fibrotic disease.

## Acknowledgements

The authors would like to thank Dr. Nicole J. Darling for assistance with drawing schematics in Figures 1 and 2. This work was supported by funding from the American Thoracic Society Research Foundation Program (CMM), the National Heart, Lung, and Blood Institute of the National Institutes of Health (NIH) under awards R01 HL153096 (CMM, TC, RB, and DWHR) and T32 HL 07085 (RB); the National Cancer Institute of the NIH under award R21 CA252172 (CMM and RB); the National Science Foundation under award 1941401 (CMM and RH); the Department of the Army under award W81XWH-20-1-0037 (CMM).

## CRediT Author Statement

**Thomas Caracena:** Methodology, Validation, Investigation, Writing – Original Draft. **Rachel Blomberg:** Formal analysis, Investigation, Writing – Original Draft, Supervision. **Rukshika S. Hewawasam:** Validation, Resources. **David W.H. Riches:** Supervision, Writing – Review & Editing. **Chelsea M. Magin:** Conceptualization, Methodology, Writing – Original Draft, Visualization, Supervision, Funding acquisition.

## Data Availability

The raw and processed data required to reproduce these findings are available here: Caracena, Thomas; Blomberg, Rachel; Hewawasam, Rukshika; Riches, David; Magin, Chelsea (2022), “Transitional alveolar epithelial cells and microenvironmental stiffness synergistically drive fibroblast activation in three-dimensional lung models”, Mendeley Data, v2 http://dx.doi.org/10.17632/58k4dh2prp.2.

