## Supplementary Materials for "Transitional alveolar epithelial cells and microenvironmental stiffness synergistically drive fibroblast activation in three-dimensional hydrogel lung models"

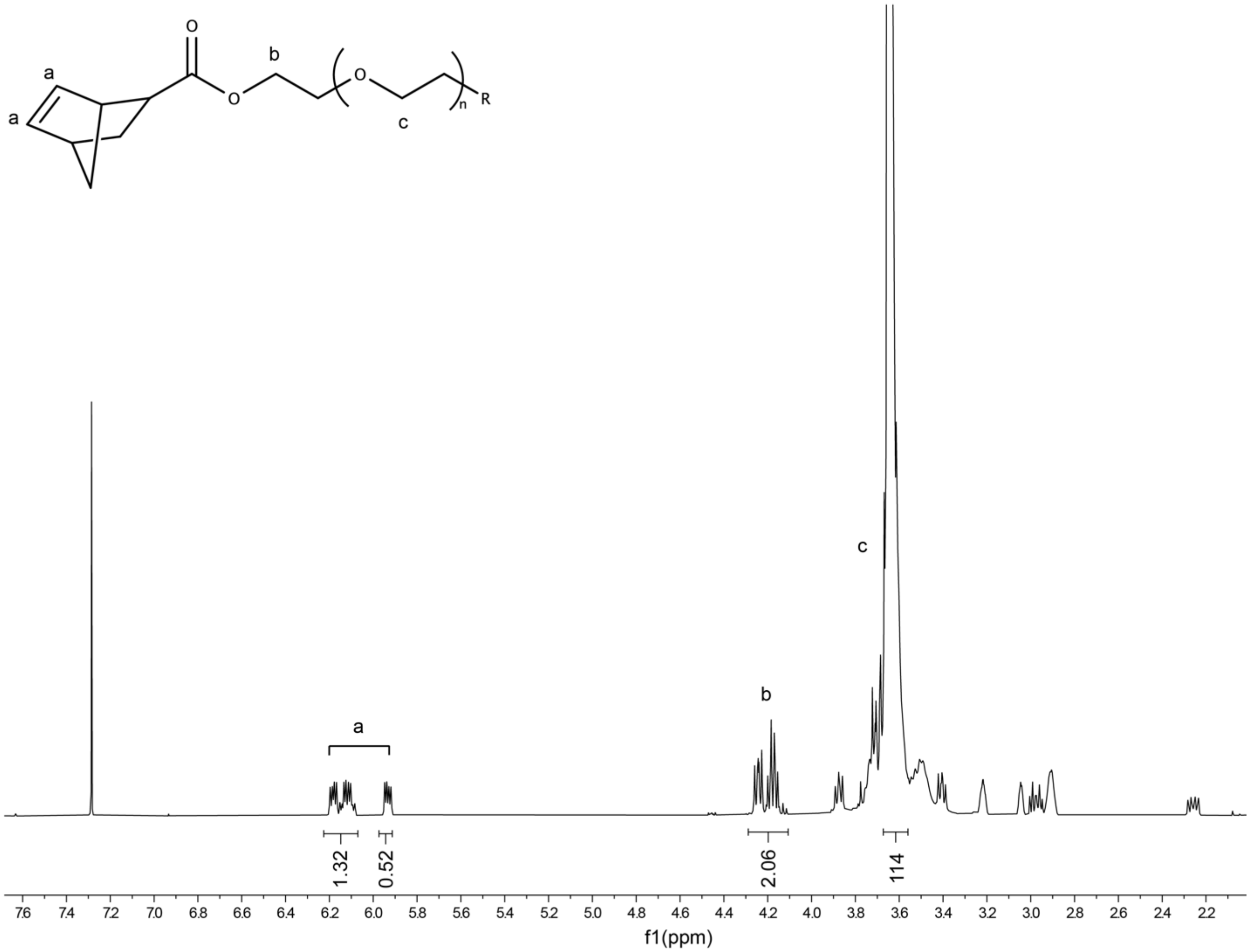


Figure S1. $^1$H NMR of 10 kg/mol poly(ethylene glycol) norbornene (PEGNB). δH (ppm) (300 MHz, $CDCl_3$, $Me_4Si$): 3.71 (s, 114H, PEG $CH_2$-$CH_2$), 4.1-4.2 (m, 2H, -$CH_2$-O), 5.9-6.2 (m, 2H, –CH=CH-). NMR shows quantitative norbornene functionalization based on a comparison of the alkene protons from norbornene to the theoretically expected number of alkene protons, using the ethylene glycol protons as a reference. Functionalization was measured as 92%.

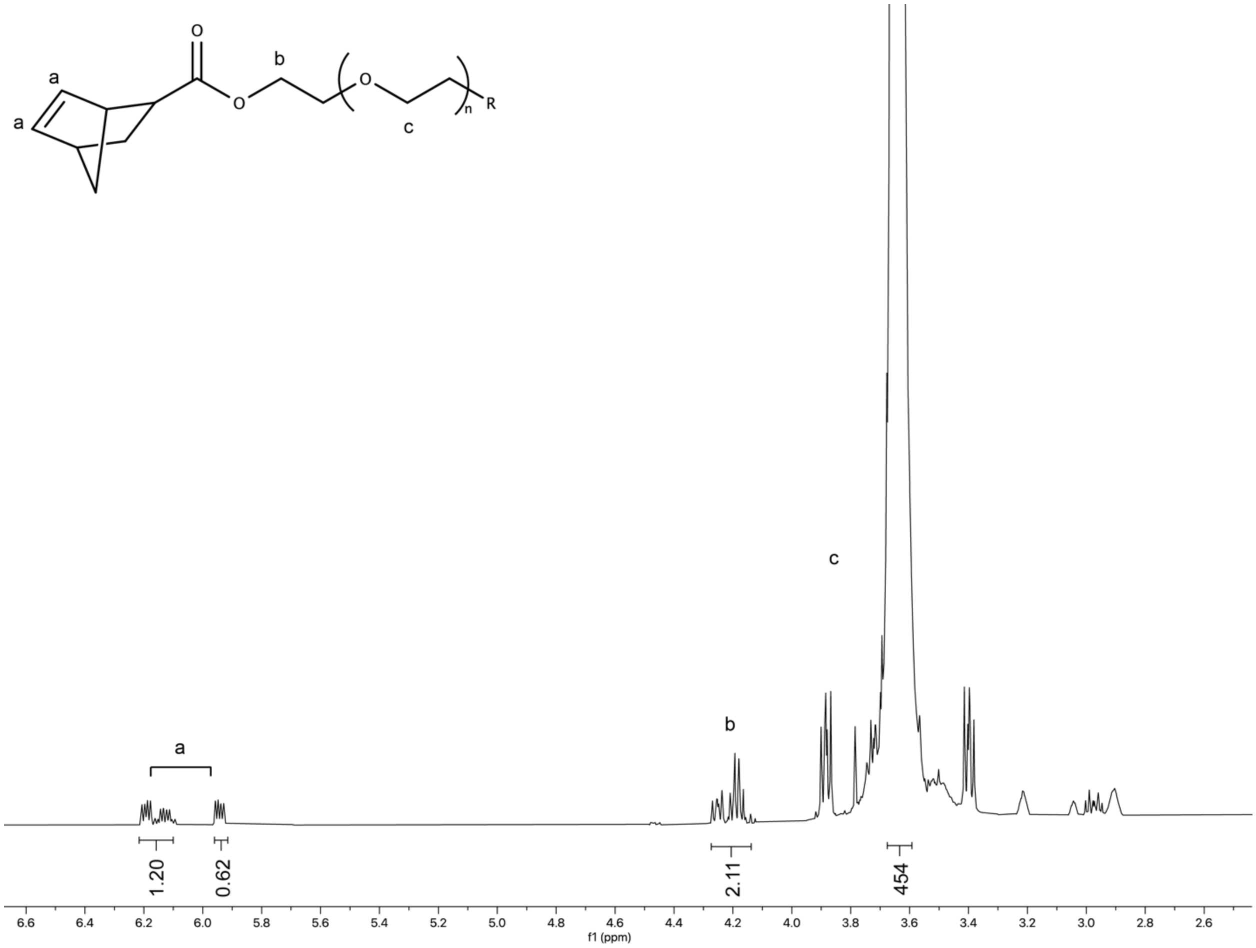


Figure S2. [1]H NMR of 40 kg/mol poly(ethylene glycol) norbornene (PEGNB). δH (ppm) (300 MHz, $CDCl_3$, $Me_4Si$): 3.71 (s, 114H, PEG $CH_2$-$CH_2$), 4.1-4.2 (m, 2H, -$CH_2$-O), 5.9-6.2 (m, 2H, –CH=CH-). NMR shows quantitative norbornene functionalization based on a comparison of the alkene protons from norbornene to the theoretically expected number of alkene protons, using the ethylene glycol protons as a reference. Functionalization was measured as 91%.
